# Low activity microstates during sleep

**DOI:** 10.1101/067892

**Authors:** Hiroyuki Miyawaki, Yazan N. Billeh, Kamran Diba

## Abstract

A better understanding of sleep requires evaluating the distinct activity patterns of the brain during sleep. We performed extracellular recordings of large populations of hippocampal region CA1 neurons in freely moving rats across sleep and waking states. Throughout non-REM (non-rapid eye movement) sleep, we observed periods of diminished oscillatory and population spiking activity lasting on the order of seconds, which we refer to as “LOW” activity sleep states. LOW states featured enhanced firing in a subset of “LOW-active” cells, and greater firing in putative interneurons compared to DOWN/OFF states. LOW activity sleep was preceded and followed by increased sharp-wave ripple (SWR) activity. We also observed decreased slow-wave activity (SWA) and sleep spindles in the hippocampus local-field potential (LFP) and neocortical electroencephalogram (EEG) upon LOW onset, but only a partial rebound immediately after LOW. LOW states demonstrated LFP, EEG, and EMG patterns consistent with sleep, but frequently transitioned into microarousals (MAs) and showed EMG and LFP spectral differences from previously described small-amplitude irregular activity (SIA) during quiet waking. Their likelihood increased over the course of sleep, particularly following REM sleep. To confirm that LOW is a brain-wide phenomenon, we analyzed data from the entorhinal cortex of rats, medial prefrontal cortex, and anterior thalamus of mice, obtained from crcns.org and found that LOW states corresponded to markedly diminished activity simultaneously in all of these regions. We propose that LOW states are an important microstate within non-REM sleep that provide respite from high-activity sleep, and may serve a restorative function.

## Introduction

The brain passes through multiple distinct stages during sleep. Each stage produces distinct activity patterns, presumably playing important roles in the function of the brain. Non-REM (non-rapid eye movement) sleep, in particular, has been associated with sleep homeostasis, synaptic plasticity, memory consolidation, and a host of other sleep functions (Rasch and Born, 2013; Rechtschaffen, 1998; Tononi and Cirelli, 2014). The signature activity pattern of non-REM sleep is the slow oscillation, an approximately 1 Hz alternation in cortical populations between UP states with robust (“ON”) spiking activity and DOWN states where most neurons are silent or “OFF” (Steriade et al., 1993). During UP/ON states, activity patterns can further display various faster oscillations, including sleep spindles (10-16 Hz) and hippocampal sharp-wave ripples (SWRs; 130 – 230 Hz) (Buzsaki, 2015). On the other hand, DOWN/OFF states are characterized by the lack of any spiking activity lasting on the order of ~100 ms (Vyazovskiy et al., 2009).

A number of sleep functions have been attributed to the slower, longer lasting microstates of the brain. For example, the transition from the DOWN to the UP state during the slow oscillation can synchronize activity across the brain and provide a window for information transfer between the hippocampus and neocortex (Rasch and Born, 2013; Sirota and Buzsaki, 2005). Meanwhile, the transition to the DOWN state could induce synaptic long-term depression (Tononi and Cirelli, 2014) in cortical networks. The DOWN state also provides neurons with respite from intensive discharge and ionic flux. It has been suggested that this respite allows for more efficient cellular restoration and maintenance (Vyazovskiy and Harris, 2013). Even longer lasting low and high activity phases have been reported during “infra-slow” oscillations under anesthesia and waking (Aladjalova, 1957; Chan et al., 2015; Filippov et al., 2008; Hiltunen et al., 2014; Lorincz et al., 2009). The high activity phases of these oscillations have been linked to “default mode” networks that subserve cognition (Chan et al., 2015; He and Raichle, 2009; Hiltunen et al., 2014), while the low activity phases reflect decreased blood flow and metabolic cost (Logothetis and Wandell, 2004).

Recently, we performed long duration recordings from the hippocampus and neocortex of freely moving and sleeping rats and report the prevalence of low firing periods that lasted several seconds, far beyond the durations attributed to DOWN/OFF states. Following Pickenhain and Klingberg (1967) and Bergmann et. al. (1987), and because of their effects on neuronal firing, we call these “LOW” activity sleep states, but they have also been called “sleep small-amplitude irregular activity” (S-SIA; Jarosiewicz et al., 2002). We reserve the term SIA, as it was originally used, to describe quiet waking patterns (Vanderwolf, 1971). We evaluate the occurrence of LOW activity sleep across the circadian cycle, and describe its effects on other oscillatory activities within non-REM sleep, including slow-wave activity (SWA), sleep spindles, and sharp-wave ripples, and on firing rates of neurons in the hippocampus and other brain regions. We contrast these with SIA states during quiet waking and microarousals (MAs), that are similar to LOW but display different electromyogram (EMG), local field potential (LFP) and electroencephalogram (EEG) patterns. We conjecture that LOW states provide neuronal rest within sleep and may prepare the brain for transitioning into quiet waking.

## Results

We performed spike-detection and unit-isolation of CA1 neurons on multiple sessions from both light and dark cycles, in four animals. In spike rasters from non-REM sleep, we observed striking and sporadic epochs of diminished activity lasting several seconds (Figure 1A-C). These epochs, which we refer to as “LOW” states, were accompanied by strongly diminished power in the LFP. To detect these low activity epochs independently of firing rates, we calculated and set thresholds on the LFP power spectrum < 50 Hz (Figure 1B, D; see Methods). Histograms of this low-passed LFP power showed two peaks in non-REM, reflecting LOW and non-LOW sleep (Figure 1B), yet only one peak in other behavioral states. We calculated a modulation index (*MI*) for each cell comparing firing rate between LOW states (*n* = 6,901) and non-REM packets (*n* = 6,411 defined as non-REM sleep excluding LOW states or MAs; see Methods). The distribution of *MI* was heterogeneous (Figure 1E and H), with a small subset of cells showing significantly positive *MI*. But overall, the mean *MI* was significantly < 0 for pyramidal cells (*MI* = −0.47 ± 0.01, *p* = 5.0 ×10-^246^, Wilcoxon signed-rank test (WSRT), multi-units (*MI* = −0.62 ± 0.01, *p* =3.3 × 10^−36^, WSRT) and interneurons (*MI* = −0.24 ± 0.02, *p* = 1.7× 10^−18^, WSRT), indicating that the balance of excitation to inhibition is shifted towards relatively higher inhibition during LOW states.

**Figure 1.**
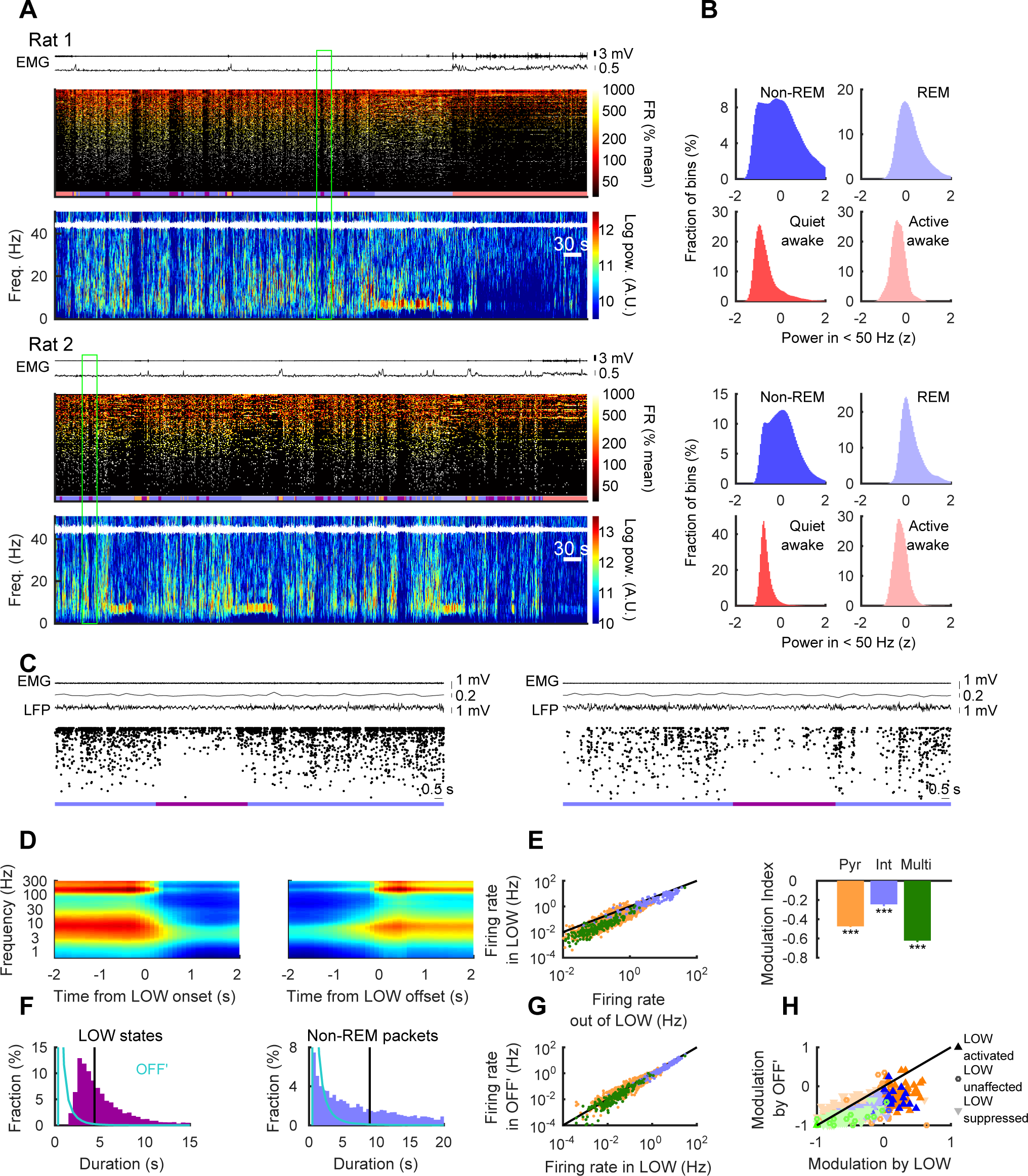
LOW activity microstates during non-REM sleep. (A) Representative recordings from hippocampal region CA1. Top panels show EMG trace (top and bottom traces depict nuchal and correlation EMG, respectively) and raster plots of firing rates in 1-s bins (179 [/108] pyramidal cells, 13 [/2] interneurons and 42 [/3] multi-unit clusters for Rat 1 [/Rat 2], ordered by mean firing rate), color normalized to the session mean. Color band beneath raster shows detected brain states (pink = quiet waking, purple/dark-blue = LOW/non-LOW states within non-REM, light blue = REM, brown = microarousals). Bottom panels show power spectrograms of hippocampal LFP (trace in white). (B) Histograms of z-scored LFP power < 50 Hz in different brain states. Non-REM histogram has two peaks, reflecting LOW and non-LOW states. (C) Zoomed examples of green boxes from panel A. (D) LOW state onset-and offset-triggered CA1 LFP power spectra (for LOWs > 2 s). (E) Firing rates of pyramidal cells (*n* = 1599, orange), interneurons (*n* = 138, blue) and multi-unit clusters (*n* = 213, green) were lower within LOW states during non-REM. Black line is identity. Right panel shows mean modulation indices (S.E.M. error bars). (F) Durations and inter-event intervals of LOW and OFF’ states. Vertical lines indicate median duration (4.4 s) and inter-event intervals (9.0 s). Distributions for OFF’ states (cyan) are superimposed (median of duration [/inter event intervals] = 450 [/400] ms). (G) Firing rates within LOW and OFF’ states were similar, by definition (colors as in panel E). (H) Most cells decreased firing within LOW states (1,438 pyramidal cells, 116 interneurons, and 191 multi-unit clusters), but a small number of “LOW-active” cells increased firing (67 pyramidal cells, 17 interneurons, and 2 multi-unit clusters) or did not change significantly (94 pyramidal cells, 5 interneurons, and 20 multi-unit clusters). Very few cells showed increased [/unchanged] firing within OFF’ (13 [/8] pyramidal cells, 1 [/0] interneuron, and 1 [/5] multi-unit clusters). ** *p* < 0.01, *** *p* < 0.001.

In these recordings, we observed right-skewed distributions for the durations of LOW states (median = 4.4 s, range: 1.5 s to 74.2s), and their inter-event intervals (median = 9.9 s; Figure 1F). The 1-s windows we used for spectral calculations put a lower limit on the duration of detectable LOW states. Nevertheless, the mode of the histogram was at 2.75 s. The distributions of these variables were also right-skewed for OFF states detected using standard methods (Johnson et al., 2010; Vyazovskiy et al., 2009), but had shorter durations (median = 0.079 s, range: 0.050-3.494 s), and inter-event intervals (median = 0.109 s). No secondary peaks were observed in the any of the histograms, indicating the absence of oscillatory cycles. On average 33.8% of LOW (*n* = 19 sessions) consisted of OFF states. However, OFF states rarely lasted > 2s (14 out of 1.2 x 10^6^ detected OFFs). An alternate method for OFF state detection (Ji and Wilson, 2007) yielded similar results (median duration = 0.070s; range: 0.010 – 4.900 s; see also Ji and Wilson’s (2007) Supplementary Figure 12d), with only 51 OFF states > 2s (out of 547,733 detected OFF states).

To further examine whether LOW states may be long-lasting OFF states, we iteratively relaxed the threshold of OFF to yield low-firing OFF’ states (Figure 1G) with the median firing rates equal to LOW states. 74.0% (*n* = 19 sessions) of LOW qualified as OFF’ states but 65.2% of detected OFF’ occurred outside of LOW. Further, 35.8% of LOW states were in long OFF’ (> 2s) while 35.1% of long OFF’ was outside of LOW. Thus, LOW and OFF’ states overlapped but were not identical. As expected, virtually all (Figure 1H, 98.7% of *n* = 1599) neurons were significantly suppressed during OFF’ states. However, a subset of cells were active specifically during LOW states (4.2%, *p* = 3.2 ×10^−30^, chi-square test; see also Figure 2C). Previous work indicates that these hippocampal “LOW-active” cells have place fields within the animal’s sleep environment (Jarosiewicz et al., 2002; Jarosiewicz and Skaggs, 2004a; Kay et al., 2016). Consistent with a recent report (Kay et al., 2016), they showed a lower firing increase during SWRs than other cells (*p* = 2.4 × 10^−6^ and 1.2 × 10^−6^, Tukey-Kramer test). Additionally, interneurons in general were more active during LOW than during OFF’ (4.2 × 10-^24^, WSRT). Overall, these observations demonstrate that LOW and OFF/DOWN states co-occur but remain distinct.

**Figure 2.**
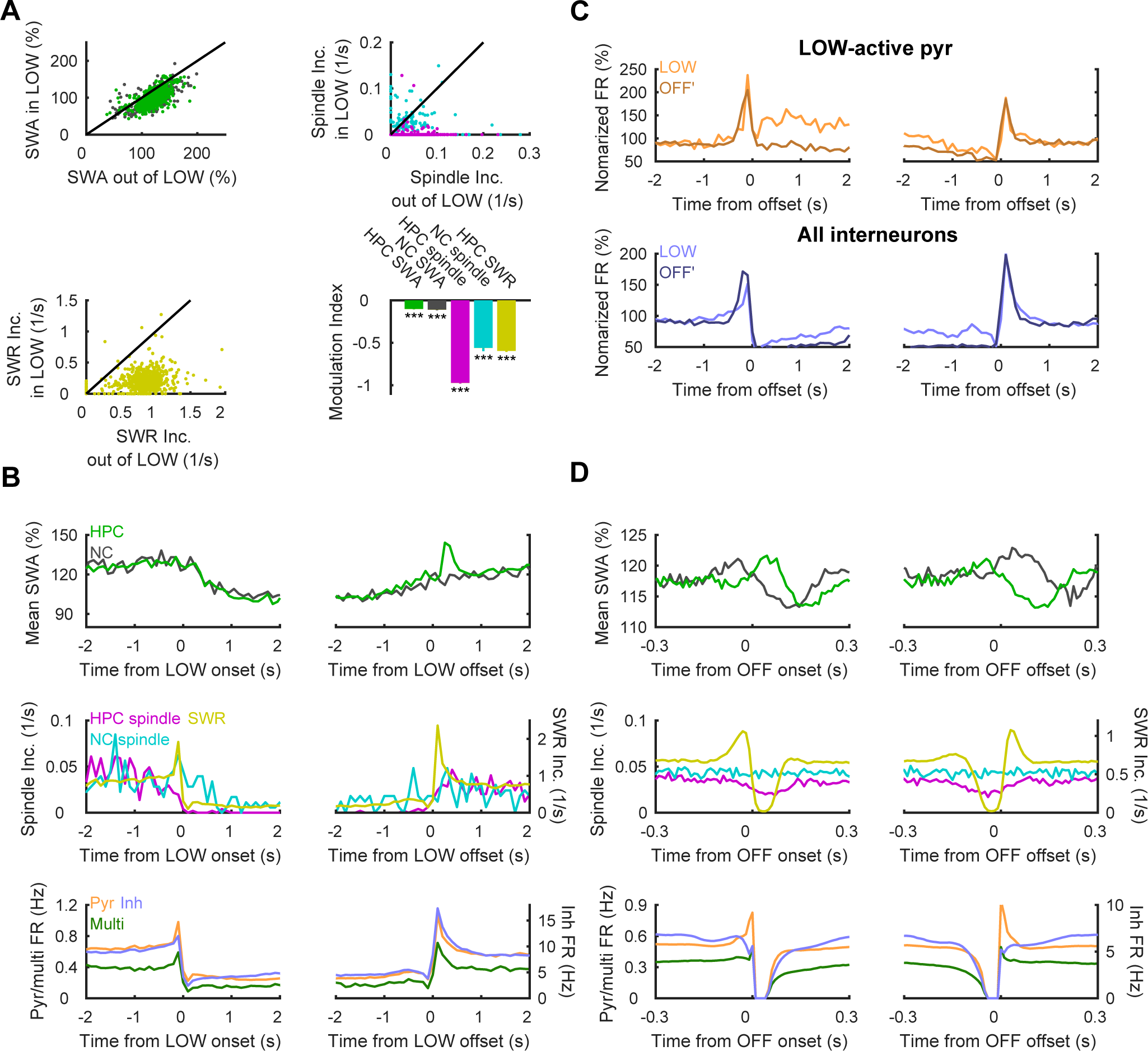
LOW state modulation of network oscillations. (A) Panels show mean slow-wave amplitudes (SWA) in hippocampus (HPC; green) and neocortex (NC; gray), the incidence rates of HPC spindles (magenta), NC spindles (cyan) and HPC SWRs (yellow) for different non-REM epochs (*n* = 927 for HPC and *n* = 436 for NC). Right panel shows the modulation indices (S.E.M. error bars). (B, C) Peri-event triggered histograms of oscillatory events and neuronal firings. (B) SWA (top) decreased after transitions to LOW states (> 2 s), and partially rebounded after transitions out of LOW states in both HPC (green; peak at 0.25 s) and NC (gray). Incidences of spindles and SWRs (middle) decreased upon LOW onset. NC spindle changes (left axis, cyan) were delayed (~ 0.5 s) relative to the HPC. LOW states were often preceded by SWRs (right axis, yellow; peak at 0.1 s) and followed by rebound SWRs (peak at 0.1s). Mean firing rates (bottom) of pyramidal cells (orange, left axis), interneurons (blue, right axis) and multi-units (green, left axis) transiently increased before onset and following offset of LOW states, likely because of SWRs. (C) Modulation of SWA (top) and spindles and SWRs incidence rates (middle), and neuronal firing (bottom) by OFF states. Note the shorter timescales compared to panel B. (D) LOW-active pyramidal cells (top, *n* = 67) showed increased firing during LOW, following a transient decrease. In contrast, the same cells showed decreased firing in OFF’ states > 2s, despite some overlap between OFF’ and LOW. Interneuron firing (bottom, *n* = 138) was higher in LOW than OFF’, although lower than non-REM. Error bars indicate S.E.M, *** *p* < 0.001.

### Oscillatory and spiking activities before and after LOW states

To examine how LOW sleep affects other oscillatory activities in non-REM, we detected slow waves (0.5– 4 Hz), sleep spindles (10–16 Hz), and sharp wave ripples (SWRs; 130–230 Hz) in the CA1 region LFP, and compared these before, during and after LOW states (Figure 2A). Not surprisingly, the mean amplitudes of hippocampal slow waves (SWA) and the incidences of hippocampal spindles and SWRs were all significantly lower during LOW. To test for more global effects of LOW states, in 8 sessions (from 2 animals), we also detected SWA and spindles on electroencephalogram (EEG) above the neocortical frontal lobe. We observed lower SWA (*MI* = −0.11 ± 0.006, *p* = 1.4 × 10^−34^) and a lower incidence of sleep spindles (*MI* = −0.56 ± 0.04, *p* = 1.6 × 10-^26^) in the neocortical EEG during LOW sleep states. We next compared activity 0.5 – 1.5s after LOW offset to 0.5 – 1.5s before LOW onset. While hippocampal SWA showed an immediate overshoot after transition out of LOW sleep (Figure 2B), SWA remained diminished in both hippocampus ( Δ*M* = −4.83 ± 1.13%, *p* = 0.006) and neocortex (Δ*M* = −9.27 ± 1.78%, *p* = 1.2 × 10^−6^). The incidence of spindles and SWRs also dropped and rose sharply at the onsets and offsets of LOW states, respectively. In particular, SWR incidence peaked 0.1 s before the transitions into and 0.1s after transitions out of LOW, indicating that SWRs tended to precede as well as follow LOW state transitions (Jarosiewicz et al., 2002). Similar to SWA, SWRs (Δ*M* = −0.178 ± 0.02 s^−1^, *p* = 4.6 × 10^−11^), hippocampal spindles (Δ*M* = −0.019 ± 0.003 s^−1^, *p* = 4.5 × 10^−7^), neocortical spindles (Δ*M* = −0.010 ± 0.006 s^−1^, *p* = 0.026), and neuronal firing rates (pyramidal cells Δ*M* = −0.085 ± 0.007 Hz, *p* = 8.7 × 10^−20^, interneurons Δ*M* = −0.45 ± 0.09 Hz, *p* = 7.8 × 10^−5^, multi-unit Δ*M* = −0.007 ± 0.027 Hz, *p* = 5.5 × 10^−4^) all remained lower for 0.5 – 1.5s after LOW offset, except in LOW-active cells (Figure 2C) and did not return to pre-LOW levels until ~10s after LOW offset (not shown). In contrast, OFF states elicited similar modulations but activities rebounded completely, and were not remain significantly different following OFF offset (Figure 2D). Overall, these analyses demonstrate that neuronal activities transiently increase immediately before and after LOW, but are strongly diminished during LOW and remain lower beyond its terminus.

### LOW sleep across cortical regions

Based on the modulation of neocortical EEG activities by LOW states, and the neocortical origin of spindles and slow waves detected in the CA1 region (Hahn et al., 2006; Isomura et al., 2006), we hypothesized that LOW states reflect a global decrease of activity throughout the cortex. To test this hypothesis directly, we analyzed two additional datasets from multiple brain regions obtained from http://crcns.org (Mizuseki et al., 2014; Peyrache and Buzsáki, 2015). Applying the same detection methods to data from the entorhinal cortex (EC) and the hippocampus of rats recorded by Mizuseki et. al. (2014), we confirmed the occurrence of LOW states in hippocampal region CA1 with coincident LOW states in layer 2/3, and layer 4/5 of the medial entorhinal cortex (Figure 3A). For the second dataset recorded by Peyrache et. al. (Peyrache and Buzsáki, 2015) in mice, we found simultaneous LOW states in the medial prefrontal cortex (mPFC), anterodorsal thalamic nucleus (ADn), post-subiculum (PoS) and hippocampus (Figure 3B and C). In both these datasets, the durations and inter-event intervals of LOW states detected in the CA1 region were distributed similarly to our original dataset (Figure 3D).

**Figure 3.**
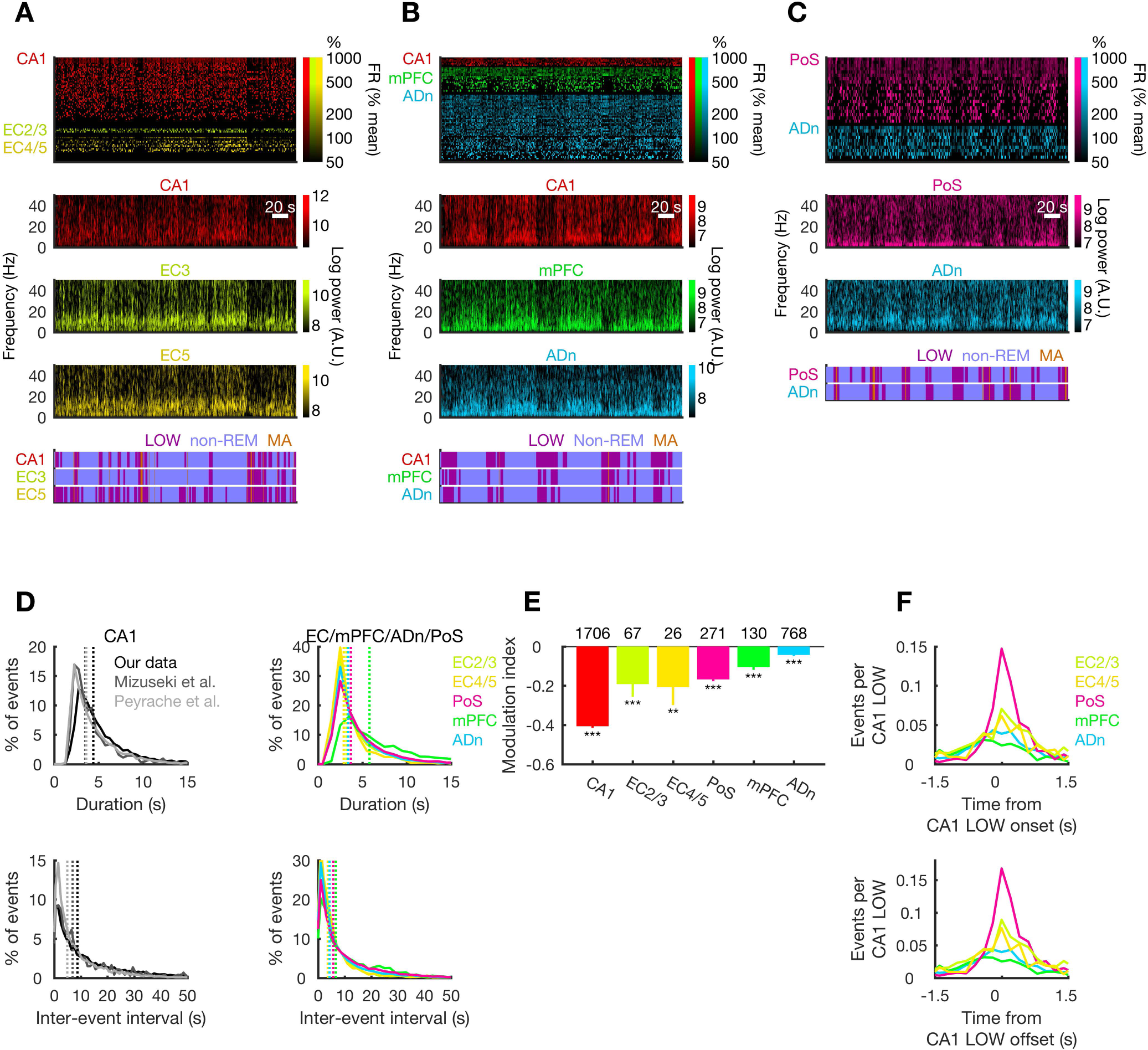
LOW states across different brain regions. (A-C) Examples of LOW states in datasets from rat (2014) (A) and mouse (2015) (B & C). Top panels show spike raster plots in 1-s bins. Color indicates brain regions, and brightness indicates firing rates normalized to the session means (46 CA1 cells, 6 EC2/3 cells, and 15 EC4/5 cells in A; 7 CA1 multi-units, 19 mPFC cells and 47 ADn cells in B; 21 PoS cells and 11 ADn cells in C). Middle panels depict power spectra of simultaneously recorded LFPs. Bottom panels show frequently synchronous LOW states detected separately for these regions. (D) Durations and inter-event intervals for LOW states in CA1 (left panels) and other brain regions (right panels) were distributed similarly across data sets and brain regions. Vertical lines indicate median values. (E) Firing rates were modulated by LOW states in all brain regions (*n* = numbers of cells, indicated above; S.E.M. error bars). (F) Cross-correlograms between LOW states in CA1 and other brain regions show largely synchronous LOW state onsets and offsets. ** *p* < 0.01, *** *p* < 0.001.

Firing rates of neurons were significantly modulated in all brain regions considered (WSRT; Figure 3E): *MI* = −0.41 ± 0.008 (*p* = 7.1 × 10^−210^) in CA1, −0.19 ± 0.064 (*p* = 7.9 × 10^−4^) in EC2/3, −0.21 ± 0.091 (*p* = 0.0010) in EC4/5, −0.17 ± 0.010 (*p* = 8.1 × 10^−36^) in PoS, −0.10 ± 0.014 (*p* = 1.1 × 10^−10^) in mPFC, −0.04 ± 0.005 (*p* = 3.3 × 10^−21^) in ADn. The strongest modulation observed in CA1 and weakest (but significant) modulation seen in ADn (*p* = 3.6 × 10^−184^, one-way ANOVA). The modulation index was significantly higher in CA1 compared to each of the other brain regions (Tukey-Kramer test, p < 0.05) and significantly lower in ADn than other regions except EC 4/5 and mPFC (Tukey-Kramer test, p < 0.05). In addition, the cross-correlogram between onsets/offsets of LOW states in CA1 and those in other regions showed clear peak around zero (Figure 3F) with no secondary peaks. Auto-correlograms of LOW onsets/offsets (not shown) also showed no secondary peaks, again indicating that LOW was not reliably cyclic. To assess the temporal propagation of LOW between brain regions, we compared onsets times within 1-s immediately before and after the CA1 LOW onset. Transition rates to LOW were higher in EC2/3, and PoS (Δ*M* = 0.045 ± 0.018 s^−1^, *p* = 0.012 for EC2/3,, Δ*M* = 0.159 ± 0.019 s^−1^, *p* = 7.4 × 10^−17^ for PoS) immediately after CA1 LOW onsets, but were greater in mPFC preceding CA1 onset (Δ*M* = −0.040 ± 0.012 s^−1^, *p* = 7.6 × 10^−4^) and unchanged in EC4/5 and ADn (Δ*M* = 0.024 ± 0.023 s^−1^, *p* = 0.30 for EC4/5, Δ*M* = 0.0004 ± 0.056 s^−1^, *p* = 0.95 for ADn). These results indicate that LOW states are global and synchronous across brain regions and propagate across cortical regions, with a large fraction initiating earlier in prefrontal regions (Massimini et al., 2004).

### Likelihood of LOW decreases within non-REM and increases across sleep

We next examined when LOW states occur. In non-REM that followed REM sleep, LOW states were present throughout the epoch (Figure 4A) but were most frequent immediately after the end of REM (Jarosiewicz et al., 2002). In non-REM epochs that terminated in waking, the likelihood of LOW states increased over time, eventually transitioning into the quiet waking state (Figure 4A). Indeed, while most (98.0%) LOW states did not terminate in waking, 62.1% of successful transitions from non-REM sleep to waking were transitions out of LOW sleep. Since LOW states were more prevalent following REM sleep and before waking, we considered that they may prepare the brain for a potential transition to waking. Consistent with this conjecture, we found a significant negative correlation between SWA (a well-established measure of sleep pressure (Achermann et al., 1993) here evaluated exclusively outside of LOW) and the fraction of time in LOW (*r* = −0.21, *p* = 2.9 × 10^−21^; Figure 4B). Moreover, we observed a lower likelihood of LOW following 3 hrs of prolonged track running (6 – 9 a.m. each day; Figure 4C), when sleep pressure was highest (Achermann et al., 1993; Miyawaki and Diba, 2016), but this likelihood increased and plateaued following sleep.

**Figure 4.**
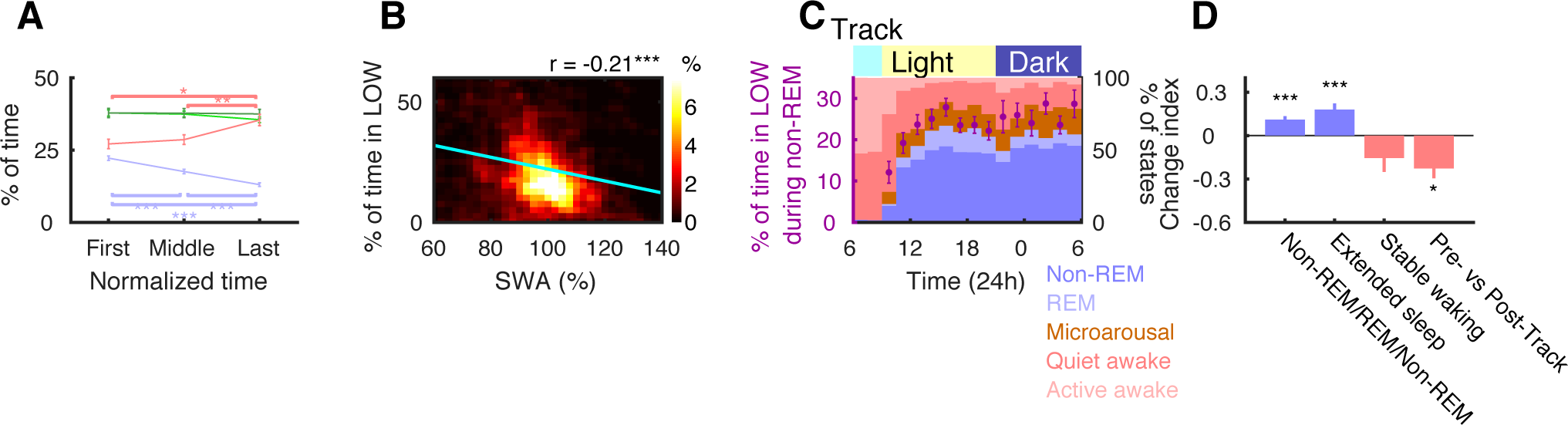
LOW states increased with decreasing sleep pressure. (A) The fraction of time in LOW decreased within non-REM epochs followed by REM (blue, *n* = 427 non-REM epochs, *p* = 9.6 × 10^−10^, one-way ANOVA followed by Tukey-Kramer tests), but increased within non-REM epochs followed by quiet waking (pink, *n* = 233 non-REM epochs, *p* = 2.6 × 10^−3^). The fraction of time in OFF states was unchanged in non-REM epochs followed by either REM (light green) or quiet waking (dark green). (B) SWA, calculated exclusively outside of LOW, was inversely correlated with the fraction of time in LOW states (*n* = 927 non-REM epochs). Regression line (cyan) is superimposed on density map of epoch means. (C) The fraction of time (within non-REM) in LOW states was lowest immediately following 3-hr track running sessions (from 6 – 9 a.m. each day), gradually increasing and stabilizing. Background colors indicates % of time spent in each behavioral/sleep state. (D) Change indices for fraction of time in LOW between non-REM epochs interleaved by REM (*n* = 295) and between the first and last non-REM epochs in extended sleep (*n* = 90) were significantly > 0. In contrast, those between non-REM epochs immediately before and after 3-hr track sessions (*n* = 7) and those between the first and last non-REM epochs in stable waking episodes (*n* = 35) decreased without reaching significance. * *p* < 0.05, ** *p* < 0.01, *** *p* < 0.001, error bars indicate S.E.M.

Likewise, the fraction of time spent in LOW states (within non-REM) increased with time (*r* = 0.086, *p* = 0.046) in extended sleep sequences of REM and non-REM (defined as sleep lasting > 30 min without interruptions > 60 s). Comparing the first and last non-REM epochs within each extended sleep, the fraction of time in LOW was significantly increased (change index *CI* = 0.18 ± 0.04, *p* = 5.9 × 10^−5^, WSRT; Figure 4D). Similarly, in non-REM/REM/non-REM triplets, the fraction of time in LOW was higher in the second non-REM (*CI* = 0.11 ± 0.02, *p* = 1.1 × 10^−8^). These changes in the frequency of LOW accounted for a large fraction of the firing rate decreases across sequential non-REM sleep epochs (*R*^2^ = 0.42, *p* = 2.5 × 10^−38^). However, sleep-dependent firing rates decreased both outside and within LOW states and previously reported correlations between firing decreases and spindles and SWRs (Miyawaki and Diba, 2016) remained significant when LOW periods were excluded (*r* = −0.30 *p* = 8.3 × 10^−8^ and *r* = −0.22 *p* = 0.0001 for spindles and SWRs, respectively). In contrast to sleep, the fraction of time in LOW decreased following the 3-hr track running sessions (Figure 4D; first hour post vs. last hour pre *CI* = −0.23 ± 0.07, *p* = 0.047, WSRT). Following stable waking periods (lasting > 15 min without interruptions > 60 s), the fraction of time in the LOW state also decreased but failed to reach significance (Figure 4D; *CI* = −0.16 ± 0.10, *p* = 0.11). These results demonstrate that LOW states are more frequent with increasing sleep, after sleep pressure has dissipated, and decrease following wakefulness, as sleep pressure accumulates.

### SIA states in quiet waking and MA

A review of the literature reveals that LOW sleep states share features with SIA and other low activity and microarousal (MA) states observed during quiet waking (Aladjalova, 1957; Chan et al., 2015; Filippov et al., 2008; Halasz et al., 2004; Hiltunen et al., 2014; Jarosiewicz et al., 2002; Lorincz et al., 2009; Vanderwolf, 1971; Watson et al., 2016). In the following (see Table 1), we separated non-REM sleep into LOW states (low EMG, low LFP), MAs (transient high EMG), and “non-REM packets” (low EMG). Quiet waking and MAs were further separated into SIA (low LFP) and non-SIA periods. We detected MA (median = 1.5 s; Figure 5A; see also Figure 1A) that interrupt non-REM sleep using EMG alone. LOW and MA states displayed distinctly different EMG levels (Figure 5B), but frequently (34.9% of the time, *n* = 17,730) MAs were preceded by LOW states and 30.2 % of MA’s transitioned into LOW states. We explicitly excluded LOW states that precede or follow MAs from all preceding analyses. We considered that longer-lasting LOW states may eventually transition into MA. However, we observed that these pre-MA LOW states were typically short, lasting (median =) 2.0 s. Post-MA LOW states lasted (median = 1.3) s.

**Figure 5.**
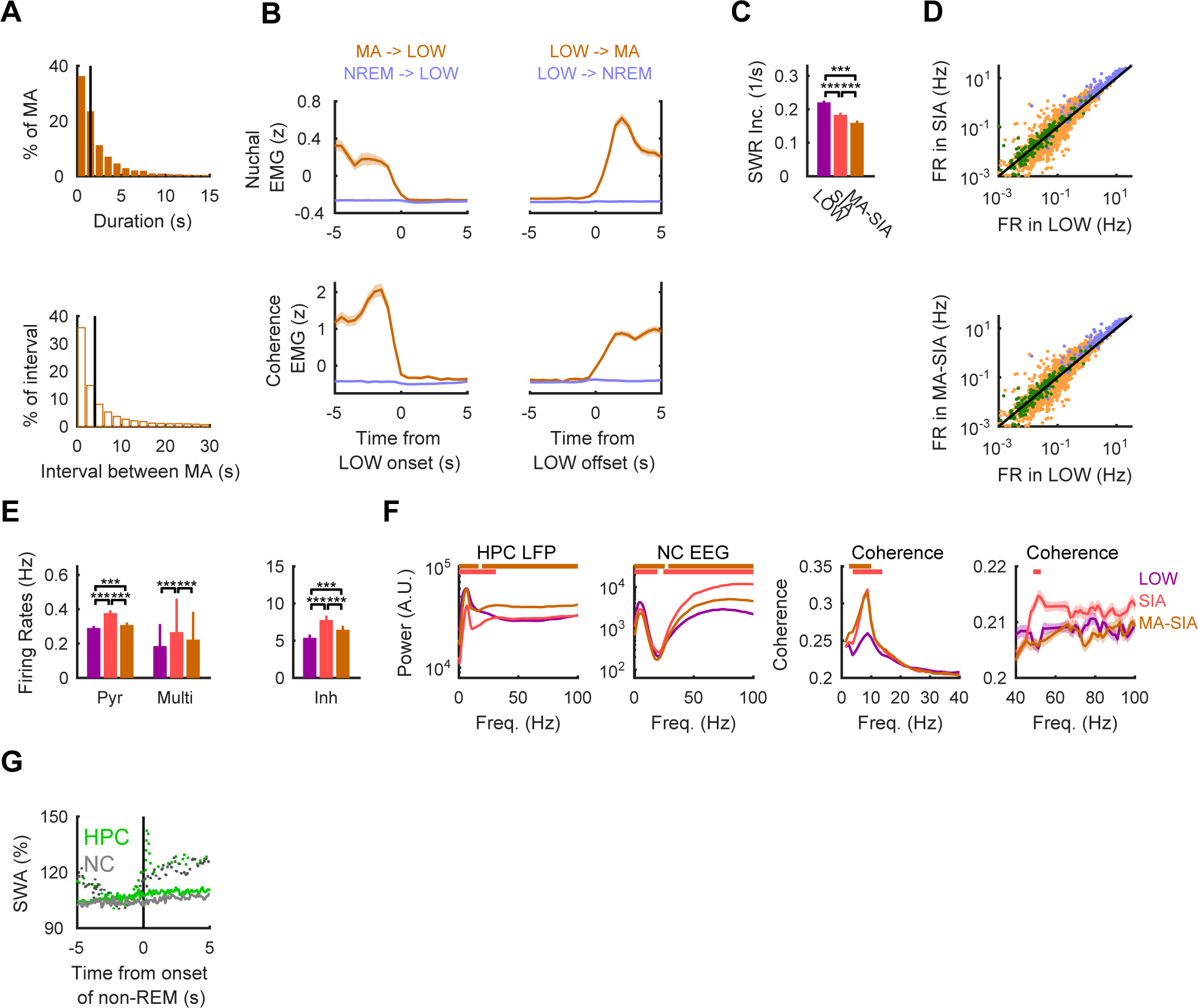
Comparison of LOW and SIA states. (A) Durations of MAs (top) and non-REM packets (NREM; bottom). Vertical lines indicate median values (1.5 s for MAs and 4.0 s for non-REM packets). (B) EMG amplitudes obtained from nuchal muscles (top) and derived from intracranial electrodes (bottom) were aligned to transitions from non-REM packets to LOW (blue) and MA (microarousal; brown). Right traces are aligned from LOW to NREM or MA. LOW transitions to MA were marked by increasing EMG amplitudes (*n* = 302 events). EMG showed virtually no change at transitions between LOW and NREM (*n* = 2151 for LOW onset and *n* = 1573 for LOW offset). Note that these analyses were limited to transition between epochs >5 s. (C) Incidence rates of SWRs in SIA and MA-SIA were lower than in LOW sleep. (D) Mean firing rates of pyramidal cells (orange), interneurons (blue) and multi-units (green) were similar in LOW and SIA. However, 23 pyramidal cells and 8 multi-unit clusters fired during LOW but not SIA (see points on x-axis; black line shows the identity). (E) Mean firing rates of pyramidal cells and interneurons were higher in SIA and MA-SIA than in LOW. (F) Power spectra of hippocampal LFP and neocortical EEG (left panels), and coherence between these regions (right panels) show significant differences in theta and gamma frequency bands (*p* < 0.001, Mann–Whitney U test with Bonferroni correction). (G) Mean SWA is compared between transitions from MA and LOW to non-REM packets (dashed lines, see Figure 2C). ** *p* < 0.01, *** *p* < 0.001, error bars indicate S.E.M. and shading indicates 95% confident intervals.

**Table 1.**
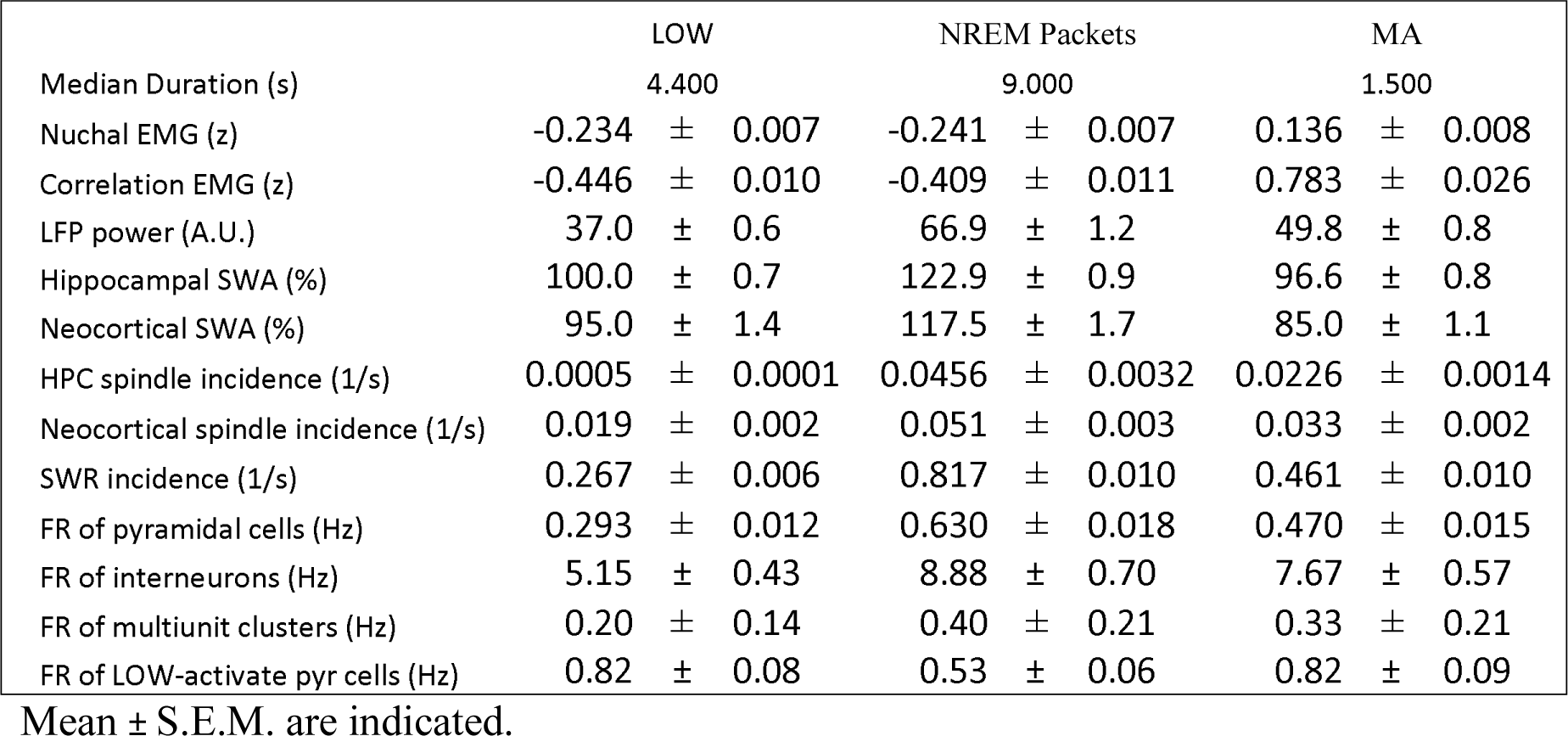

We then applied LOW detection to both MA and quiet waking periods in our recordings to see if similar activities patterns are also observed then. Following the earlier literature (Jarosiewicz and Skaggs, 2004a, b; Kay et al., 2016; Vanderwolf, 1971), we use the term “SIA” to refer to periods with low LFP power < 50 Hz (similar to LOW) during waking and MA, when EMG is high. Thus, if EMG is high 41.3 ± 2.7 % (*n* = 19) of time in MA was classified as SIA (median MA-SIA duration = 2.9 s). Additionally, much of quiet waking similarly qualified as SIA (56.1 ± 3.4 %), compared to 21.6 ± 2.6 % of non-REM in LOW. Durations and inter-event intervals of SIA states in quiet waking were distributed similarly to sleep LOW, though SIA states were slightly longer (median = 6.3 s, *p* = 2.4 × 10^−176^ Mann–Whitney U tests) with shorter inter-event intervals (median = 4.1 s, *p* = 9.3 × 10^−174^). Unexpectedly, SIA featured faster (but still small) head speed compared to other (non-SIA) quiet waking, suggestive of muscle twitches or grooming (*M* = 2.4 cm/s during SIA and *M* = 1.6 cm/s during quiet waking, *p* < 10^−300^, WSRT), but substantially less than active waking (*M* = 4.6 cm/s, *p* < 10^−300^, WSRT).

Next, we compared neuronal activity between LOW and SIA. Slow waves and spindles were not observed during waking or SIA, but SWRs were. As expected (Jarosiewicz et al., 2002), SWR incidence rates were significantly modulated by SIA (*MI* = −0.41 ± 0.017, *p* = 3.3 × 10^−88^, WSRT) and by MA-SIA (−0.61 ± 0.012, *p* = 6.6 × 10^−241^). SWR incidences in SIA and MA-SIA were lower than in non-REM LOW (Figure 5C). Firing rates of neurons were generally similar in LOW and SIA (Figure 5D). However, some units appeared to be active only during LOW, without firing in SIA (i.e. see 31 points along x-axis of Figure 5B). These cells had no reliable place fields within the home cage (in 29 out of 31 peak place field firing was < 0.3 Hz). Nevertheless, overall firing rates were slightly higher in SIA than LOW (*p* = 6.3 × 10^−18^ for pyramidal cells, 4.0 × 10^−6^ for multi-unit cluster, and 9.4 ×10^−23^ for interneurons; Figure 5E).

To further examine differences between LOW and SIA, we also compared the power spectra of the hippocampal LFP and neocortical EEG between MA-SIA, SIA and LOW states (Figure 5F). While the same threshold was used to detect both LOW and SIA, overall patterns in SIA were quite different from LOW. Hippocampal LFP had significantly lower power < 31.1 Hz in SIA states than in LOW, consistent with less synchronization during waking states, and in the neocortical EEG spectrum, SIA had slightly lower power in < 19.5 Hz and stronger power in > 24.4 Hz (Figure 5F). Moreover, hippocampus LFP and neocortical EEG demonstrated enhanced coherence near theta and gamma bands during SIA indicating a greater level of consciousness and arousal during SIA, and in theta (but not gamma) during MA-SIA, perhaps indicative of drowsiness. Therefore, SIA and LOW demonstrate Finally, SWA activity showed an immediate partial rebound following transition from LOW to non-REM (Figures 2F & 5G), but not from MA to non-REM. Thus LOW is a microstate *within* non-REM sleep, whereas MA is an *interruption* of sleep and resets the buildup of SWA (Achermann et al., 1993).

## Discussion

In summary, we observed long lasting suppressed activity periods during non-REM sleep, which we call LOW activity sleep. In earlier studies in natural sleep, microstates were observed that share features with the LOW activity sleep states we describe (e.g. Bergmann et al., 1987; Pickenhain and Klingberg, 1967; Roldan et al., 1963; Terzano et al., 1985). However, population spiking activity was not available in those studies and it could not be determined whether these low-amplitude epochs were desynchronized because of high or low neuronal firing activity. Jarosiewicz and colleagues (Jarosiewicz et al., 2002; Jarosiewicz and Skaggs, 2004a) also studied SIA and LOW in large-scale hippocampal unit recordings and made similar observations about neuronal firing and SWRs. Our study provides a quantitative confirmation and extends on their reports in multiple ways.

LOW sleep was characterized by a strong decrease in population spiking and oscillatory activities including slow waves, sleep spindles, and SWRs (Table 1). While most activities rebounded, their levels remained diminished for several seconds, indicating a lingering effect of LOW states beyond their termini (Halasz et al., 2004; Watson et al., 2016). SWRs, however, were transiently increased immediately before and after LOW, potentially reflecting homeostatic rebound (Girardeau et al., 2014), and a vulnerability for epilepsy (Penttonen et al., 1999; Vanhatalo et al., 2004), though we saw no evidence of pathology. We also showed that LOW states strongly suppress neuronal activity not just in the hippocampus, but throughout the brain, including the entorhinal cortex, prefrontal cortex, post subiculum and anterodorsal thalamus. This suppressed activity lasted much longer than typical OFF states and was accompanied by increased firing in a LOW-active subset of neurons (Jarosiewicz and Skaggs, 2004a; Kay et al., 2016). Interestingly, both OFF and LOW states were preceded by transient increased firing, suggesting a common onset mechanism, such as a hyperpolarizing current (Compte et al., 2003; Lorincz et al., 2009). But importantly, LOW states featured higher levels of interneuron firing than did OFF’ states. Ultimately, intracellular recording are needed to test whether DOWN level membrane potentials (Steriade et al., 1993) are present during LOW and further examine commonalities and differences between LOW and DOWN/OFF states.

LOW sleep occurred not just after REM but throughout non-REM sleep, and may be triggered in response to high neuronal activity, from either REM sleep or UP/ON states and SWRs (Miyawaki and Diba, 2016; Vyazovskiy et al., 2009), that increase the need for neuronal rest and restoration. Vyazovskiy and Harris (2013) recently proposed that a global quiescent state may be valuable for cellular repair and restoration of organelles. LOW states may be suitable for carrying out such functions. We uncovered an inverse relationship between the rate (fraction of time) of LOW sleep and sleep pressure, with low likelihood in early non-REM sleep following waking, but increasing likelihood in late sleep. The neuronal activities that occur in non-REM outside of LOW states (e.g. slow waves, spindles, and SWRs) have been linked to some of the memory benefits of sleep (Buzsaki, 2015; Rasch and Born, 2013; Tononi and Cirelli, 2014) and are more prevalent in early sleep (Miyawaki and Diba, 2016). These observation, combined with the greater fidelity of replay during early sleep (Kudrimoti et al., 1999), may support the idea that early sleep serves for circuit modification, including synaptic downscaling (Tononi and Cirelli, 2014), while late sleep, under decreased sleep pressure, may provide for cellular restoration and upkeep (Genzel et al., 2014; Vyazovskiy and Harris, 2013).

We demonstrated that LOW states are distinctly a sleep state, with lower EMG, lower gamma and gamma coherence, but higher low-frequency (delta and theta) activity, and greater SWA rebound than SIA or quiet waking. Unfortunately, some confusion may arise from inconsistent terminology in the literature. For example, LOW sleep states were called “S-SIA” by Jarosiewicz et. al. (2002), but later “SIA” by the same authors (Jarosiewicz and Skaggs, 2004a, b) and more recently, by Kay et. al. (2016). They also appear to share features with phase B of the cyclic alternating pattern (CAP) in non-REM sleep (Halasz et al., 2004; Terzano et al., 1985). We propose that SIA should be reserved for the waking state, as originally described by Vanderwolf (1971) and later employed by others (Jarosiewicz et al., 2002), while LOW amplitude/LOW activity sleep is best reserved for the sleep state, as also originally used (Bergmann et al., 1987; Pickenhain and Klingberg, 1967). Nevertheless, it is feasible that collectively LOW sleep and waking SIA represent the sleep and waking extremes of a continuum of microarousals (Halasz, 1998; Watson et al., 2016). Consistent with this notion, we observed frequent transitions between LOW states and MA, and MAs demonstrated both SIA and non-SIA periods. SIA periods (relative to LOW) showed less slow activity, and higher gamma power and gamma coherence between hippocampal LFP and neocortical EEG, consistent with consciousness, and hippocampal place-cells that encode the animal’s sleep location begin to fire during both LOW sleep and SIA (Jarosiewicz et al., 2002; Jarosiewicz and Skaggs, 2004a; Kay et al., 2016), as if in preparation for spatial cognition.

LOW states also appear to share features with the low-activity phase of infra-slow oscillations (Aladjalova, 1957; Chan et al., 2015; Filippov et al., 2008; Hiltunen et al., 2014; Lorincz et al., 2009), though we observed no evidence for repeated oscillatory cycles in LOW (Figure 1E). Recent evidence suggests the low-activity phases of infra-slow oscillations are produced by long-lasting hyperpolarizing potassium currents, mediated by ATP-derived adenosine released from astrocytes (Lorincz et al., 2009). Such non-synaptic currents may produce LOW states, potentially accounting for the parallel firing decreases observed in both pyramidal cells and interneurons, as well as the lingering effect from LOW states on subsequent activity patterns (see also ref(Watson et al., 2016)). Interestingly, infra-slow low-activity phases coincide with large-scale calcium waves in astrocytic networks (Kuga et al., 2011), which may account for their global nature. Cerebral blood flow during these low-activity phases is ~10% lower, indicating that they are detectable in global BOLD activity as well as LFP (He and Raichle, 2009). Given these facts, along with the central role that astrocytes and adenosine play in cellular repair (Chen and Swanson, 2003), energy metabolism (Porkka-Heiskanen and Kalinchuk, 2011), and mediating sleep pressure (Halassa et al., 2009), they may also play a central role in the genesis and function of LOW states.

## Experimental Procedures

We analyzed data from hippocampal region CA1 with neocortical EEG described in a previous study (Miyawaki and Diba, 2016), along with data from CA1 and entorhinal cortex (Mizuseki et al., 2014), and from anterodorsal thalamus, and post-subiculum and medial prefrontal cortex (Peyrache and Buzsáki, 2015), freely available at http://crcns.org. Additional details of methods, including surgery, unit clustering, local field spectrum analysis, sleep state, slow-wave, spindle, and sharp-wave ripple detections, are provided elsewhere (Miyawaki and Diba, 2016).

### Animals, surgery, and data collection

Four male Long-Evans rats (260-370g; Charles River Laboratories, Wilmington, MA) were anesthetized with isoflurane and implanted with silicon microprobes in the dorsal hippocampus (2.00 mm lateral and 3.36 mm posterior from the bregma). In three of the rats, 2 stainless steel wires (AS 636, Cooner wire, Chatsworth, CA) were placed into the nuchal muscles to measure the electromyograph (EMG). Electrical coherence in microprobe electrodes in the 300-600 Hz band was also used as an alternate measure of EMG (Schomburg et al., 2014; Watson et al., 2016) that does not depend on nuchal electrode placement. We took filtered signals from the top channel of each shank and calculated mean pair-wise correlations among electrode pairs separated ≥ 400 µm in 0.5-s bins. These two measures were independent but strongly correlated (r = 0.63 in 5-s time bins). In two of the rats, screws were inserted on the skull above the right frontal lobe (3.00 mm lateral and 2.50 mm anterior from the bregma) and attached to Nyleze insulated copper wires (overall diameter 0.13 mm; Alpha Wire, Elizabeth, NJ). The signals were recorded with a Digital Lynx recording system (Neuralynx, Bozeman, MT), then processed using previously described methods (Diba and Buzsaki, 2008)

After a recovery period from surgery (> 5 days), rats were placed on a water-restriction regimen to motivate track running, and were given ad-libitum water for 30 mins each day. Three out of four rats were maintained in a home cage except during the last 3 h of dark cycles, during which they were put on an I-, L-, or U-shaped linear track in the same room and given water rewards on platforms after every traversal. A fourth rat was recorded only in the home cage during the light cycle. Two colored light-emitting diodes were mounted on the headstage and the animals’ movements were tracked with an overhead video camera. All procedures were in accordance with the National Institutes of Health guidelines and approved by the University of Wisconsin-Milwaukee Institutional Animal Care and Use Committee.

### Unit clustering and cell classification

Unit clustering and cell classification were performed as previously described (Diba and Buzsaki, 2008). We analyzed light cycles and dark cycles separately. For track-running sessions, we concatenated data from behavior on the track with the last 3 hrs of the preceding dark cycle (pre-sleep) and the first 3 hrs of the following light cycle (post-sleep). Putative pyramidal cells and interneurons were separated based on standard methods using spike waveforms, burstiness, refractory periods, and firing rates (Bartho et al., 2004; Csicsvari et al., 1998; Sirota et al., 2008). Units with an isolation distance < 15 (Schmitzer-Torbert et al., 2005) were considered potentially multi-units.

### Spectral analyses

LFP, EMG and EEG traces were low-pass filtered at 1250 or 1280 Hz using NDManager and its plugins (Hazan et al., 2006)(http://ndmanager.sourceforge.net). Power spectra were whitened and calculated using multitaper methods and the Chronux toolbox for Matlab (Bokil et al., 2010) in 1-s windows.

### Summary of microstate classification and terminology

Non-REM sleep was separated into LOW states (low EMG, low LFP), MAs (transient high EMG), and “non-REM packets” (low EMG) representing the remainder of non-REM sleep. Quiet waking and microarousals (MAs) were further separated into SIA (low LFP) and non-SIA. REM was characterized by low EMG and high theta, while active waking featured high EMG and high theta. Further details are provided below.

### Microstate detection

Sleep and waking were separated based on nuchal EMG and the animal’s movement. In one rat which did not have nuchal EMG, we used coherence EMG instead. Data from this animal was consistent with the rest, but was excluded from detailed EMG analyses in Figures 5B. Sleep was detected by low EMG power and no movement (defined below). The remainder was considered waking. EMG signals were first smoothed with a 1-s Gaussian filter and power was z-scored in 500-ms overlapping windows at 100-ms steps. A two-threshold “Schmitt” trigger was used to detect transitions between “low” and “high” EMG power at 0 and 0.5 threshold z scores. Similarly, the thresholds for “no movement” and “movement” were set at 0.5 cm/s and 5 cm/s. Transient (< 10 s) low EMG within waking was ignored. Transient high EMG power epochs (> 0.1 s) within sleep were marked as MAs. Detected states underwent post-hoc visual inspection and occasional manual modification. REM was inferred from high theta (described below) with no movement sandwiched between non-REM epochs. The theta (5-10 Hz) over (1-4 Hz plus 10-14 Hz) band ratio of the power spectral density was used to detect transitions between high theta and low theta, using custom-made MATLAB software written by Anton Sirota (Sirota et al., 2008) based on the Hidden Markov Model Toolbox for Matlab (Kevin Murphy), followed by visual inspection. Sleep states with high theta were classified as REM (rapid eye movement) and the remainder were classified as non-REM (Grosmark et al., 2012; Robinson et al., 1977). Similarly, waking periods with high theta were labeled “active awake” and the remainder were labeled “quiet waking.” For the Peyrache et al. (2015) dataset, we used the provided non-REM, REM, and wakefulness timestamps, and considered only epochs > 50 sec.

### LOW/SIA activity detection

LOW states were detected based on the power spectra of the LFP in each brain region calculated in 1-s windows sliding with 0.1-s steps. The average power between 0 – 50 Hz was Gaussian filtered (σ = 0.5 s) and z-scored based on mean and SD within non-REM sleep. Histogram of filtered power during non-REM had two peaks (Figure 1B). We determined the position of lower peak and the local minimum between the two peaks based on second derivative of smoothed (Gaussian filter, σ = 0.1 z-score) histogram. Periods in which z-scored power was lower than the local minimum were detected as candidate LOW states. If the LFP power in a candidate LOW did not drop below the lower peak of the histogram, it was discarded. Two consecutive LOW states separated by < 0.5 s were concatenated. LOW states that even partially coincided with MAs were excluded from Figure 1–3 analyses. We used these same thresholds to detect SIA during quiet waking and MA.

For analyses in Figure 2, which required more accurate timestamps, onsets and offsets were determined from population (MUA) firing rates. First, the mean firing rate in previously detected LOW and bordering non-LOW intervals were calculated. The threshold for each LOW onset/offset was set to the mean firing rate in LOW plus 20% of the difference with bordering non-LOWs and onset and offset timestamps were shifted accordingly.

### Sharp-wave ripple detection

SWRs were detected following previously described methods (Diba et al., 2014). First, the ripple band (130-230Hz) power of the LFP was calculated during non-REM and quiet waking. Channels with the largest power in the ripple band were selected for each shank, and periods with power exceeding 1 SD of the mean at least one of the selected channels were labeled as candidate events. Candidates with short gaps (< 50ms) were combined. Candidates shorter than 30 ms or longer than 450 ms were abandoned. Candidates were classified as SWRs if their peak power was greater than 5 SDs of the mean.

### Sleep spindle detection

Hippocampal LFP and neocortical EEG were band-pass filtered between 10–16 Hz (nearly identical results were observed with a wider 9–18 Hz band (Miyawaki and Diba, 2016)). For hippocampal spindles, the channel with the largest mean power during non-REM sleep was used. Candidate spindles in each signal were detected when amplitudes of the Hilbert transform exceeded 1.5 SDs above the mean for longer than 350 ms (Sullivan et al., 2014). Candidates were concatenated when inter-event intervals were shorter than 125 ms. Candidates with peak amplitudes below 4 SDs above the mean were abandoned. For each candidate, spindle troughs were detected. Candidates with inter-trough intervals longer than 125 ms (corresponding to slower than 8 Hz) were discarded.

### Slow wave detection

We used a previously established method (Miyawaki and Diba, 2016; Vyazovskiy et al., 2011; Vyazovskiy et al., 2009) to detect individual slow waves in the neocortical EEG and the hippocampal LFP as positive deflections between two negative deflections in the band-pass filtered signal (0.5 – 4 Hz). The polarity of the hippocampal LFP was inverted due to the polarity reversal relative to the EEG (Nir et al., 2011). Slow waves separated by less than 100-ms were discarded. Slow wave activity (SWA) was defined by the amplitude of these individual slow waves measured from trough to peak.

### OFF/OFF’ state detection

Hippocampal OFF states arise from neocortical influence during SWA (Hahn et al., 2012; Hahn et al., 2006; Isomura et al., 2006). OFF states during non-REM were defined as periods with no CA1 units spiking for ≥ 50 ms (Grosmark et al., 2012; Johnson et al., 2010; Luczak et al., 2007; Vyazovskiy et al., 2009). To test whether LOW states might be equivalent to long-lasting OFF states under relaxed assumptions, we detected low-firing periods (OFF’ states) by thresholding Gaussian filtered (σ = 100 ms) multi-unit activity (MUA) in 50-ms bins within non-REM sleep (Ji and Wilson, 2007). The threshold for each session was set at the value for which the median MUA was equivalent within OFF’ and LOW states.

### Modulation/Change indices

LOW modulation index (MI) for variable X was defined as (X_LOW_ – X_non-REM packets)_ / (X_LOW_ + X _non-REM packets_) for LOW and (X_SIA_ – X_non-SIA_) / (X_SIA_ + X _non-SIA_) for SIA. The change index (CI) for X was defined by (X_post_ – X_pre_)/(X_post_ + X_pre_).

### LOW/OFF’ modulated cells

Spikes were counted in 100-ms bins, then means within LOW and OFF’ states were calculated. The same number of bins were then randomly selected from outside of LOW, to calculate surrogate means. This procedure was iterated 2000 times within each session separately. If the mean within LOW/OFF’ was higher (or lower) than top (or bottom) 0.5% of the shuffled data, the cell marked as activated (or suppressed). We obtained qualitatively similar results with a novel community detection method (Billeh et al., 2014).

## Acknowledgements

We are grateful to Adrien Peyrache, Kenji Mizuseki, and crcns.org for making their data readily available, and Christof Koch, Markus Schmidt, and Brendon O. Watson for valuable comments.

## Author Contributions

H.M. performed research. Y.N.B. performed analyses identifying LOW-active cells.

H.M. and K.D. designed research and wrote the manuscript

